# GPX4 Promotes Optic Nerve Regeneration and RGC Neuroprotection

**DOI:** 10.1101/2025.06.03.657677

**Authors:** Ming Yang, Fuyun Bian, Xue Feng, Liang Li, Haoliang Huang, Liang Liu, Roopa Dalal, Hang Yang, Pranav Varma Suraparaju, Frank Cao, Petrina Ong, Alexandria Luo, Dong Liu, Yang Hu

## Abstract

Preventing retinal ganglion cells (RGCs)’s soma and axon degeneration and promoting optic nerve (ON) regeneration holds great promise for effective glaucoma treatment. To explore potential neural repair strategies, we focused on glutathione peroxidase 4 (GPX4), a key regulator of lipid peroxidation. GPX4 is upregulated in surviving RGCs after acute ON crush or chronic ocular hypertension insult, and also in regenerating RGCs. AAV-mediated RGC-specific overexpression of GPX4 promotes significant ON regeneration and RGC survival, along with visual functional preservation, demonstrating the detrimental role of lipid peroxidation in glaucoma and the therapeutic potential of modulating lipid peroxidation through GPX4 in optic neuropathies.

## INTRODUCTION

Glaucoma is recognized as the leading cause of irreversible blindness globally, projected to affect over 100 million individuals by 2040^1^. The progressive degeneration of retinal ganglion cells (RGCs) and their axons in the optic nerve (ON) is the hallmark of glaucomatous neurodegeneration^2-4^. Therefore, preventing RGC and ON degeneration and/or promoting ON regeneration holds great promise for advancing glaucoma treatment.

Recently, lipid peroxidation and ferroptosis have been linked to experimental mouse models of optic neuropathies^5-7^. Iron-dependent, excessive phospholipid peroxidation leads to membrane destruction and subsequently, a regulated cell death named ferroptosis that is different from apoptosis or necroptosis^8,9^. Ferroptosis can be suppressed by iron chelation/depletion or lipophilic radical trapping/reduction. Glutathione peroxidase 4 (GPX4) is the major reductase of phospholipid hydroperoxides (PLOOHs), a lipid-based reactive oxygen species (ROS), acting with a tripeptide (cysteine, glycine, and glutamic acid) cofactor glutathione (GSH). Loss of GPX4 induces lipid peroxidation-dependent ferroptosis^10^.

In this study, we found GPX4 upregulation in surviving RGCs after ON crush (ONC) or ocular hypertension injuries, and also in regenerating RGCs. RGC-specific overexpression of GPX4 promotes significant ON regeneration and RGC survival, demonstrating the role of lipid peroxidation in neurodegeneration and the neural repair potentials of modulating lipid peroxidation through GPX4 in optic neuropathies.

## RESULTS

### GPX4 is upregulated in crushed and regenerating RGCs

Naïve mouse RGCs have a modest expression of GPX4, which is not changed much 7 days post ON crush (7dpc) but significantly increased in the surviving RGCs at 14dpc (**Fig. 1A**), when the majority of injured RGCs are dead, indicating the neuroprotective role of GPX4. The expression pattern of GPX4 protein in crushed RGCs is reminiscent of GPX4 transcript dynamic changes published before using single-cell RNA-seq^11^ (**Fig. S1**). Interestingly, the GPX4 transcript is also upregulated in regenerating RGCs (regRGCs) (**Fig. 1B**) detected by Retro-Seq analysis, a retrograde tracing-sequencing scheme we developed before to differentially label, purify, and sequence regRGCs and non-regenerating surviving RGCs (surRGCs)^12^, suggesting its pro-regeneration activity.

**Figure 1.**
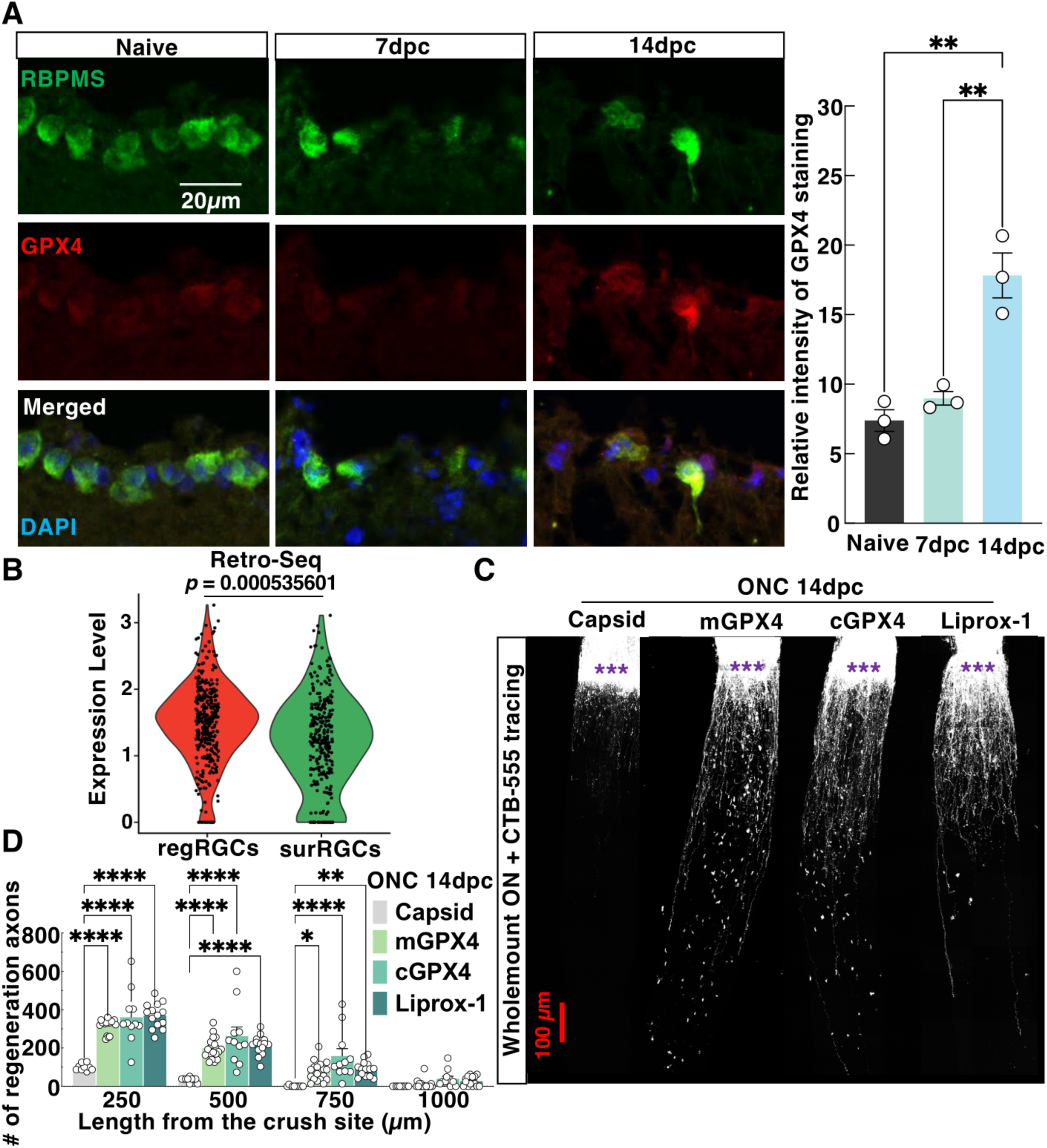
GPX4 is upregulated in surviving and regenerating RGCs, which promotes significant ON regeneration after ONC injury. (**A**). Representative confocal images of retina sections with immunostaining of RGC marker RBPMS and GPX4. The GPX4 fluorescent intensity is quantified in naïve and ONC mice at 7-day post-optic nerve crush (7dpc) and 14dpc. Scale bar: 20 µm. n=3 retinas/group. Data are presented as means ± s.e.m. **: *p* < 0.01, one-way ANOVA with Tukey’s multiple comparison tests. (**B**). GPX4 mRNA levels in regenerating and non-regenerating surviving RGCs (regRGCs *vs* surRGCs) were detected by Retro-seq. *p* = 0.000535601. Violin plots show the distribution of GPX4 expression in individual cells, with each dot representing a single cell. Wilcoxon Rank Sum test, and the p-value was adjusted for multiple comparisons. (**C**). Confocal images of ON wholemount after optical clearance showing maximum intensity projection of regenerating fibers labeled with CTB-Alexa 555 at 14dpc. Scale bar, 100 μm. ***: Crush site. (**D**). Quantification of regenerating fibers at different distances distal to the lesion site. Data are presented as means ± s.e.m, n=10 of AAV-Capsid group, n=18 of AAV-mGPX4 group, n=11 of AAV-cGPX4 group, n=14 of Liprox-1 group. Data are presented as means ± s.e.m, *: *p* < 0.05, **: *p* < 0.01, ****: *p* < 0.0001, two-way ANOVA with Tukey’s multiple comparisons test.

### RGC-specific GPX4 overexpression promotes ON regeneration after crush injury

There are three isoforms of GPX4 in mammalian cells: mitochondrial GPX4 (mGPX4), cytosolic GPX4 (cGPX4), and nucleolar GPX4 (nGPX4). They share the same amino acid sequence at the C-terminus, including the enzymatic active site, but differ at the N-terminus, which contains signal sequences that direct their subcelluar distributions^13^. GPX4 is a selenoprotein whose translation is contingent upon the presence of the Selenocysteine Insertion Sequence (SECIS), a conserved RNA element located in the 3′ UTR that is critical for the incorporation of selenocysteine at an in-frame UGA codon, which would otherwise serve as a termination signal during translation. We focused on mGPX4 and cGPX4 and engineered AAV constructs to drive their expression with the RGC-specific mSncg promoter^14^ and the respective SECIS elements to facilitate accurate selenocysteine incorporation (**Figure S2A**). We then confirmed RGC-expression of GPX4 after AAV intravitreal injection: HA-tagged GPX4 colocalized with RBPMS^+^ RGCs (**Fig. S2B**). We next examined the effect of RGC-specific GPX4 overexpression on ON regeneration after ONC in mice *in vivo*. Both forms of GPX4 induced significant ON regeneration although cGPX4 generated slightly more and longer regenerating axons than mGPX4 (**Fig. 1C,D**). Our result indicates that inhibition of lipid peroxidation may promote ON regeneration. Lipophilic radical-trapping antioxidant, liproxstatin-1 (Liprox-1), is a potent small molecule inhibitor of lipid peroxidation^10,15^, which is preferred to be used in vivo due to its enhanced efficacy, improved chemical stability, and favorable pharmacokinetic properties^9,16^. Liprox-1 acts as radical-trapping antioxidant^15^ to decrease PLOOHs, similar to GPX4 function (**Fig. S2C**). Consistently, we found that Liprox-1 treatment via intraperitoneal injection also induced significant ON regeneration (**Fig. 1C,D**). In additon to lipid peroxidation, iron is required for ferroptosis. However, to our surprise, administration of ferroptosis inhibiting iron chelator deferiprone (DFP) did not induce significant ON regeneration, although it increased RGC survival after ONC (**Figure S3**), suggesting the lack of involvement of iron-dependent ferroptosis in axon regeneration. These findings collectively suggest that the inhibition of lipid peroxidation but not ferroptosis significantly promotes ON regeneration in the mouse ONC model, highlighting the potential therapeutic benefits of targeting lipid peroxidation for neural repair.

### ONC-induced lipid peroxidation in RGC death

To determine lipid peroxidation directly, we examined two end-products of lipid peroxidation in RGCs: malondialdehyde (MDA)^17^ and 4-Hydroxynonenal (4-HNE)^18^. We confirmed that ONC injury induced elevation of MDA in ONs (**Fig. 2A**) and of 4-HNE in ganglion cell layer (GCL) (**Fig. 2B,C**). Consistently, GPX4 treatment significantly decreased MDA and 4-HNE (**Fig. 2A-C**). Inhibiting lipid peroxidation by mGPX4, cGPX4, and Liprox-1 all increased RGC survival after ONC (**Fig. 2D,E**). Together with significant RGC neuroprotection by DFP (**Fig. S3D,E**), our results indicate that both lipid peroxidation and ferroptosis contribute to ON traumatic injury-induced RGC death.

**Figure 2.**
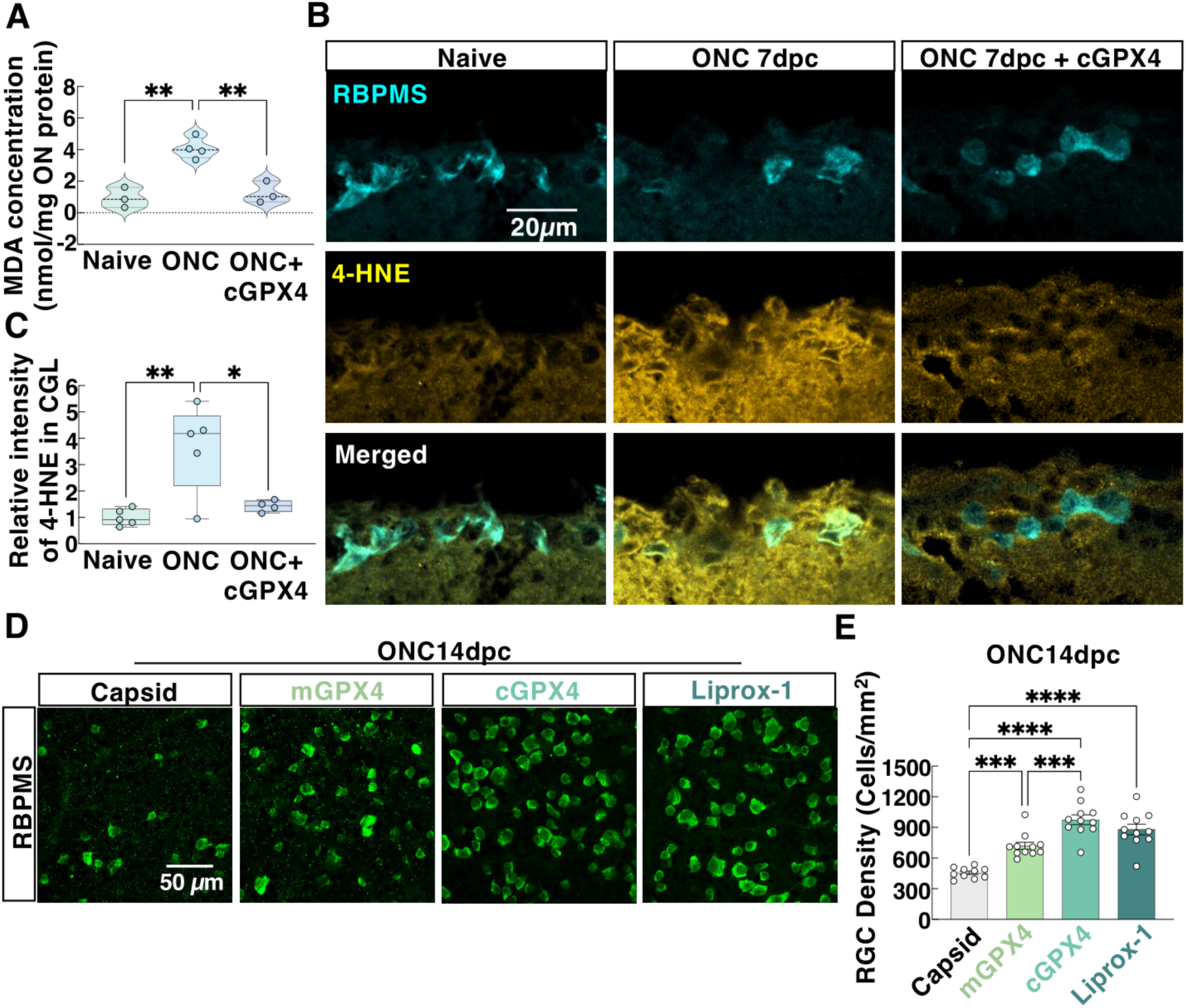
GPX4 decreases RGC lipid peroxidation and promotes RGC survival after ONC. (**A**) MDA levels of ON in naïve and ONC mice at 7dpc were determined (each dot represents one sample pooled from 4 ONs). Naïve (n=3, 12 ONs), ONC (n=4, 16 ONs), ONC+AAV-cGPX4 (n=3, 12 ONs). (**B**) 4-HNE was label by immunostaining in retina section of naïve, ONC, and ONC+cGPX4 mice at 7dpc. Scale bar, 20 µm. (**C**) Quantification of relative fluorescent intensity of 4-HNE in the retinal ganglion cell layer (GCL). Naïve (n=5 retinas), ONC (n=5 retinas), ONC+AAV-cGPX4 (n=4 retinas). (**D**) Representative confocal images of the flat-mounted retinas at the peripheral regions showing surviving RBPMS-positive RGCs at 14dpc. Scale bar, 50 µm. (**E**): Quantification of surviving RGC somata from peripheral flat-mounted retinas, AAV-Capsid (n=10), AAV-mGPX4 (n=11), AAV-cGPX4 (n=11), Liprox-1 (n=11). (**A,C,E**) Data are presented as means ± s.e.m, *: *p* < 0.05, **: *p* < 0.01, ***: *p* < 0.001, ****: *p* < 0.0001, one-way ANOVA with Tukey’s multiple comparisons test.

### GPX4 prevents glaucomatous neurodegeneration and preserves visual functions in the mouse glaucoma model

We also found GPX4 elevation in glaucomatous RGCs by RiboIP-seq^19^ (**Fig. 3A**) and confirmed by immunostaining (**Fig. 3B**). RGC-specific overexpression of GPX4 did not affect IOP elevation in the mouse SOHU glaucoma model (**Fig. 3C**) which enables us to evaluate its RGC autonomous effect in glaucomatous neuroprotection. First, we performed *in vivo* imaging and visual function assays to evaluate GPX4’s effect in glaucoma mice^19-22^. *In vivo* OCT imaging showed significant thinning of the retinal ganglion cell complex (GCC), including retinal nerve fiber layer (RNFL), GCL and inner plexiform layer (IPL), in glaucoma eyes injected with control AAVs compared to contralateral naïve control eyes, at 3 weeks post SO injection (3wpi) (**Fig. 3D,E**), indicating significant glaucomatous neurodegeneration. GPX4 treatments significantly increased GCC thickness, again, slight better with cGPX4 than mGPX4 (**Fig. 3D,E**). Moreover, *in vivo* assessment of RGC electrophysiologic function by pattern electroretinogram (PERG) and of visual acuity by optokinetic tracking response (OKR) demonstrated that GPX4 treatments significantly preserved visual function of the glaucoma eyes (**Fig. 3F,G**). We next investigated RGC somata and axons survival by histological analysis of postmortem retina wholemounts and ON semi-thin sections. Quantification of surviving RGC somata in retina and surviving RGC axons in ONs consistently showed that GPX4 treatments strikingly increased RGC survival throughout the peripheral, middle, and central regions of the glaucomatous retinas and RGC axon survival of the glaucomatous ONs (**Fig. 3H-L**). However, ferroptosis inhibitor DFP treatment did not dramatically improve visual function in glaucoma mice (**Fig. S4A-D**), although it slightly increased GCC thickness and RGC and ON survival (**Fig. S4E-J**). Taken all together, these findings suggest that the inhibition of lipid peroxidation significantly promotes glaucomatous neuroprotection.

**Figure 3.**
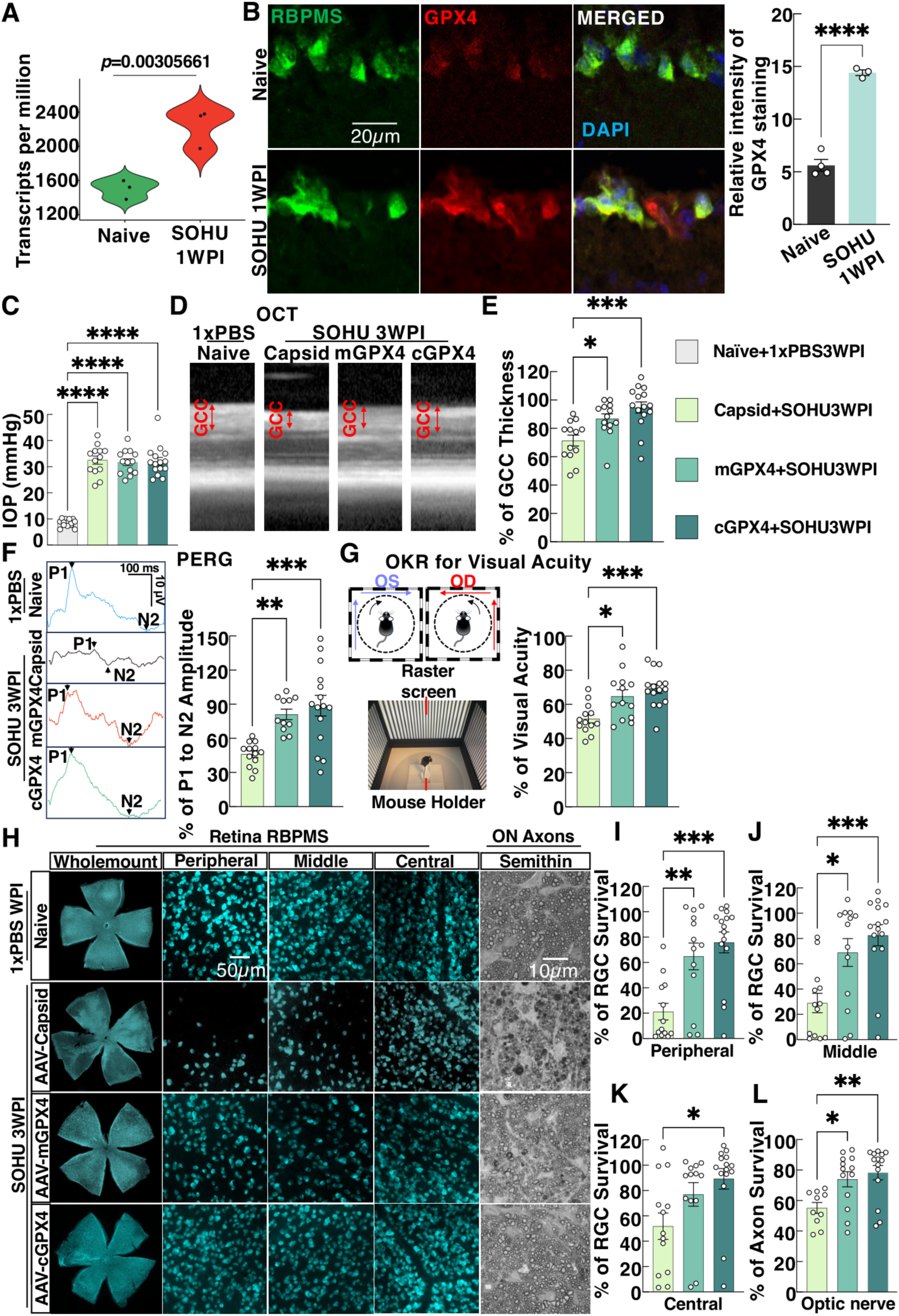
GPX4 promotes significant neuroprotection and preserves visual function in the SOHU glaucoma model. (**A**) GPX4 mRNA levels detected by bulk RGC RiboTag-Seq, *p* = 0.00305661. Violin plot showed the distribution of GPX4 transcripts per million (TPM) in each group, n=3 per group, with the Wald test. (**B**) Representative confocal images of retina sections with immunostaining of RGC marker RBPMS and GPX4. The GPX4 fluorescent intensity is quantified in naïve and glaucoma mice at 1wpi. Scale bar: 20 µm. n=3 retinas/group. Data are presented as means ± s.e.m., ****: p < 0.0001, Student’s t-test. (**C**) IOP of naïve and glaucomatous eyes with or without AAV2-GPX4 treatments at 3wpi. Naive mice with intracameral injection of 1xPBS as naïve control (n=14), AAV-Capsid injected SOHU mice (n=13 mice), AAV-mGPX4 injected SOHU mice (n=13), and AAV-cGPX4 injected SOHU mice (n=15 mice). (**D**) Representative in vivo OCT images of mouse retinas at 3WPI. GCC: ganglion cell complex, including RNFL, GCL, and IPL layers; indicated as double-end arrows. (**E**) Quantification of GCC thickness was measured by OCT at 3WPI, represented as the percentage of GCC thickness in the SOHU eyes compared to the contralateral control eyes. (**F**) Left, representative waveforms of PERG. Right, quantification of P1-N2 amplitude of PERG at 3WPI, represented as a percentage of glaucomatous eyes compared to the contralateral control eyes. AAV-Capsid (n=13 mice), AAV-mGPX4 (n=11 mice), AAV-cGPX4 (n=15 mice). (**G**) Visual acuity measurement by optomotor, represented as the percentage of glaucomatous eyes compared to the contralateral control eyes. (**H**) Representative images of the whole flat-mounted retinas showing surviving RBPMS-positive RGCs at 3WPI and images of peripheral, middle, and central flat-mounted retinas; scale bar: 50µm. The far-right panel: representative light microscope images of ON semi-thin transverse sections with paraphenylenediamine staining at 3WPI. Scale bar: 10µm. (**I,J,K**) Quantification of surviving RBPMS+ RGCs at peripheral, middle, and central retinas, represented as a percentage of glaucomatous eyes compared to the contralateral control eyes. (**L**) Quantification of surviving axons in ON semithin sections, represented as the percentage of glaucomatous eyes compared with the contralateral control eyes. AAV-Capsid (n=10 mice), AAV-mGPX4 (n=13 mice), AAV-cGPX4 (n=14 mice). All the quantification data of C,E,F,G,I, J, K, L are presented as means ± s.e.m. *: *p* < 0.05, **: *p* < 0.01, ***: *p* < 0.001, ****: *p* < 0.0001, one-way ANOVA with Tukey’s multiple comparison tests.

## DISCUSSION

Here, we demonstrated the significant role of lipid peroxidation in RGC axon regeneration and RGC survival, blocking of which by PLOOH reductase GPX4 or lipophilic radical-trapping antioxidant Liprox-1 achieved ON regeneration and RGC neuroprotection. Surprisingly, ferroptosis inhibitor/iron chelator DFP increases RGC survival dramatically in the ONC mouse model and modestly in the glaucoma model, it does not promote ON regeneration, indicating distinct roles of lipid peroxidation and ferroptosis on CNS axon regeneration. Although neuron survival is not invariably linked proportionately with axon regeneration^23^, lipid peroxidation and ferroptosis have been inevitably linked together^8-10^. It is very intriguing to reveal the molecular mechanism responsible for the different effects of blocking lipid peroxidation and ferroptosis on axon regeneration, which may lead to a deeper understanding of how axon regeneration happens and how ferroptosis results from lipid peroxidation. One attractive idea is that plasma membrane structure and associated membrane/submembrane signaling molecules are critical for axon regeneration, based on our previous studies^12^. Lipid peroxidation is highly enriched in the cell membrane, which leads to decreased diffusivity and membrane fluidity^24^. The current result shed more light on the local membrane molecules in axon regrowth: lipid condition on plasma membrane is responsible for the structure and mechanical behaviours of membranes that can contribute to membrane extension directly; oxidative stress on membrane lipid after axon injury may decrease in the bending rigidity to restrict the membrane morphological changes, thus to prevent axon regrowth. In addition, lipid rafts on the membrane are the signaling hubs that are critical for many physiological functions. Lipid peroxidation also disturbs many aspects of membrane lipid rafts, such as their stability and abundance, and protein and lipid composition^25^. These disruptions may also contribute to the neurodegeneration and the failure of axon regeneration. More research needs to be done to profile the content of lipid species on axon membrane and their redox conditions before and after axon injury under the conditions with or without axon regeneration. One step further, our study brings up an even more general possibility: the inhibitory role of oxidative stress including lipid and protein redox conditions in axon regeneration and neuroprotection.

Ferroptosis has been suggested to be involved in glaucomatous neurodegeneration^5-7^. It was recommended that both antioxidants and iron chelators need to show protection to confirm ferroptosis is in fact involved^9^. However, two of these studies only used lipid ROS scavenger ferrostatin-1^5,7^, which essentially demonstrated that lipid peroxidation is involved in glaucomatous neurodegeneration. One study indeed used iron chelator DFP but not lipid ROS scavenger to show the neuroprotective effect in pathologically high IOP-induced acute RGC death^6^. We used the same DFP in the SOHU glaucoma model and only found minimal neuroprotection, drastic different to the effect of GPX4 treatment, indicating lipid peroxidation per se but not ferroptosis is the driving force of glaucomatous neurodegeneration.

In summary, we found lipid peroxidation is induced by ON injury, which contributes to neurodegeneration and restriction on axon regeneration. The critical redox enzyme GPX4 of PLOOH can effectively block neurodegeneration and promote ON regeneration, indicating a novel therapeutic strategy on conquering optic neuropathies.

## MATERIALS AND METHODS

### Animals and Housing Conditions

Wild-type (WT) male and female C57BL/6J mice were obtained from Jackson Laboratory (Bar Harbor, ME, USA). The animals were housed under standard laboratory conditions, which included a 12-hour light/dark cycle and ad libitum access to food and water, with room temperature at 25 ± 2 °C and humidity between 40% and 60%, and both male and female mice were used for the experiments. All experimental procedures were conducted in compliance with institutional ethical guidelines and received approval from the Institutional Animal Care and Use Committee (IACUC) at Stanford University School of Medicine.

### Constructs and AAV production

The coding sequences (CDS) for MitoGPX4 (mGPX4) and CytoGPX4 (cGPX4), along with their respective selenocysteine insertion sequence (SECIS), were amplified from mouse tissue cDNA libraries using Q5 High-Fidelity DNA Polymerase (New England Biolabs, M0491L). The PCR sequences of the primers are as follows. mGPX4 Forward: MfeI-ATCGCAATTGGCCACCATGAGCTGGGGCCGTCTGAG; Reverse: XbaI-CGATTCTAGAGAGATAGCACGGCAGGTCCTTC. SECIS element for mGPX4 Forward: MluI-ACGTACGCGTTAATCTAGCCCTACAAGTGTGTGCCCCTA; mGPX4 Reverse: HindIII-CGATAAGCTTCACATTTCTACATTTTATTCCCACA. cGPX4 Forward: MfeI-ATCGCAATTGGCCACCATGTGTGCATCCCGCGATGATTGGC; cGPX4 Reverse: XbaI-ACTGTCTAGAGAGATAGCACGGCAGGTCCTTC. SECIS element for cGPX4 Forward: MluI-ACGTACGCGTTAATCTAGCCCTACAAGTGTGTGCCCCTA; SECIS element for cGPX4 Reverse:

HindIII-CGATAAGCTTCACATTTCTACATTTTATTCCCACA. The resulting amplified fragments were subsequently inserted into the vector backbones pAM-AAV-mSncg-3HA-WPRE utilizing restriction enzyme sites MfeI, XbaI, MluI, or HindIII. Plasmid constructs were purified using the Endo-Free Plasmid Maxi Kit (Omega Bio-tek, D6926-03) in accordance with the manufacturer’s instructions. Adeno-associated virus (AAV) particles were produced following the protocol outlined by Wang et al^14^. In brief, HEK293T cells were co-transfected with the AAV plasmids containing the target constructs, the pAAV2 (pACG2)-RC triple mutant (Y444, 500, 730F), and the pHelper plasmid (Stratagene) using the PolyJet transfection reagent (SignaGen Laboratories, SL100688). After 72 hours of incubation, the cells were harvested and lysed, and the AAV particles were purified through two successive cesium chloride density gradient centrifugations. Viral titers were quantified using real-time PCR and adjusted to a concentration of 1.5 × 10¹² vector genomes (vg)/mL. A total of 2 μL of the AAV solution was administered intravitreally into each eye.

### Ophthalmological procedures and measurements

The comprehensive methodologies for intravitreal injection, intraocular pressure (IOP) measurement, histological analysis of retinal ganglion cells (RGCs) and optic nerve (ON), in vivo optical coherence tomography (OCT) imaging, pattern electroretinography (PERG), and optokinetic response (OKR) have been previously published^12,14,19,20,26,27^. Brief descriptions are presented below.

### Intravitreal Injection

Mice were anesthetized using a combination of xylazine (0.01 mg/g) and ketamine (0.08 mg/g), with dosages calculated based on body weight. For the delivery of adeno-associated virus (AAV) via intravitreal injection, a finely pulled and polished glass microcapillary pipette was carefully inserted into the peripheral retina near the ora serrata of approximately 4-week-old mice. Prior to the injection, approximately 2 μL of vitreous humor was aspirated to create space for the injection of 2 μL of AAV suspension into the vitreous. Following AAV administration, the mice were bred for an additional two weeks to allow for adequate expression of the transgene. To facilitate anterograde labeling of regenerating axons, 2 μL of cholera toxin subunit B conjugated to Alexa Fluor 555 (CTB555, Invitrogen) at a concentration of 2 mg/mL was injected intravitreally.

### Compound treatments

In experiments involving pharmacological intervention with Liproxstatin-1 (Liprox-1, MedChemExpress, HY-12726), wild-type mice received intraperitoneal injections of Liprox-1 at a dosage of 10 mg/kg^10^. Treatment commenced on the day of optic nerve crush (ONC) and continued every other day for a total duration of 14 days. For deferiprone (DFP), an iron chelator treatment, mice was subjected to 14 days pre-treatment before ONC, and 14 days post-treatment of 1mg/ml DFP in drinking water. In SOHU glaucoma model, mice received 14 days of pre-treatment before the SO injection, followed by 21 days of post-treatment.

### Optic nerve crush (ONC) surgery and treatment

The ONC procedure was conducted two weeks post-administration of adeno-associated virus (AAV), when the mice were aged 6 to 7 weeks. Under aseptic surgical conditions, the optic nerve was carefully exposed intraorbitally at the 12 o’clock position, without any potential damage to the retro-orbital sinus. A standardized crush was performed to the optic nerve at approximately 0.5 mm posterior to the globe, utilizing jeweler’s forceps (Dumont #5; Fine Science Tools, Foster City, CA) for a precise duration of 5 seconds. Following the procedure, a topical neomycin-containing ophthalmic ointment (Akorn, Somerset, NJ) was applied to the cornea to provide protection and facilitate recovery.

### Assessment of malondialdehyde (MDA) level

Optic nerve MDA level was measured with an Lipid Peroxidation (MDA) Assay Kit (ab118970) following the manufacturer’s instructions. 4 optic nerves from the same group were pooled into one sample, with 3 biological replicates in each groups to ensure enough amount of protein concentration. MDA concentration was determined according to the standard curve and normalized with the corresponding protein concentrations.

### Silicon oil-induced ocular hypertention (SOHU) glaucoma model and intraocular pressure (IOP) measurement

The experimental procedures for the SOHU glaucoma model were adapted from previously established protocols^19,20,28^. Nine-week-old mice were anesthetized via intraperitoneal injection of Avertin at a dosage of 0.3 mg/g body weight. A 32G needle was employed to meticulously create a tunnel through the corneal layers at the superotemporal limbus, ensuring that the lens and iris remained undamaged, thereby facilitating access to the anterior chamber. Subsequently, a sterile, handmade glass micropipette was utilized to inject 2 μL of silicone oil (1,000 mPa·s; Silikon, Alcon Laboratories, Fort Worth, TX) into the anterior chamber. The silicone oil droplet was calibrated to expand and cover a substantial portion of the iris, achieving a diameter ranging from 1.8 to 2.2 mm. Following the injection, a veterinary antibiotic ointment (BNP Ophthalmic Ointment, Vetropolycin, Dechra, Overland Park, KS) was applied to the treated eye to mitigate the risk of infection. The contralateral eye served as a control and received a 2 μL injection of normal saline into the anterior chamber. Throughout the procedure, artificial tears (Systane Ultra Lubricant Eye Drops, Alcon Laboratories, Fort Worth, TX) were administered as necessary to maintain corneal hydration.

Intraocular pressure (IOP) measurements were conducted in both eyes prior to the silicone oil injection and at three weeks post-injection (wpi) using a TonoLab tonometer (Colonial Medical Supply, Espoo, Finland), in accordance with the manufacturer’s instructions. Mice were anesthetized with xylazine (0.01 mg/g) and ketamine (0.08 mg/g) based on body weight. To facilitate pupil dilation, 1% Tropicamide ophthalmic solution (Akorn, Somerset, NJ) was administered to the treated eyes three times at three-minute intervals, achieving full dilation within 10 minutes. IOP measurements were obtained by taking six individual readings per eye, which were averaged to generate a single machine-derived value. Three readings generated by the tonometer were recorded for the eyes, and the mean of these values was utilized to calculate the final IOP. Artificial tears were continuously dropped to the eyes to ensure corneal hydration throughout the IOP measurement procedure.

### Spectral-domain optical coherence tomography (SD-OCT) imaging

Optical coherence tomography (OCT) imaging was conducted using a 30° licensed lens (Heidelberg Engineering) in accordance with previously established protocols^20,29^. The mouse retina was scanned in central scan mode, with an average of 100 frames acquired under high-resolution settings. Heidelberg software was used for the measurement of the ganglion cell complex (GCC), which includes the retinal nerve fiber layer (RNFL), ganglion cell layer (GCL), and inner plexiform layer (IPL). The average GCC thickness surrounding the optic nerve head in the diseased retina was compared with that of the contralateral control retina to calculate the GCC thickness (%). All measurements were performed by investigators who were blinded to the sample treatments to ensure objectivity.

### Pattern electroretinogram (PERG) measurement

Simultaneous PERG recordings were conducted using the Miami PERG system (Intelligent Hearing Systems, Miami, FL), based on protocols established in our previous studies^20,29^. To maintain a consistent core body temperature throughout the procedure, the animals were placed on a heating pad with feedback-control function (TCAT-2LV, Physitemp Instruments Inc., Clifton, New Jersey), which was set to maintain a temperature of 37°C. Prior to the recordings, a small amount of lubricant eye drops (Systane) was applied to each eye to prevent the formation of corneal opacities. Electrode placement was performed as follows: the mouse was inserted with the reference electrode subcutaneously at the back of the head, positioned between the ears. The ground electrode was inserted at the base of the tail; and an active steel needle electrode was inserted subcutaneously on the snout to facilitate the acquisition of responses from both eyes simultaneously. Two LED-based stimulators, each measuring 14 × 14 cm, were positioned directly in front of the eyes, with the center of each screen located 10 cm from the respective eye. The stimulus pattern consisted of black-gray elements (spatial frequency: 0.052 cycles/degree), maintained at 85% contrast and a luminance of 800 cd/m². Upon visual stimulation, PERG signals were independently recorded from the snout using asynchronous binocular acquisition. Each trace recorded lasted up to 1020 ms, and the final readout was derived by averaging two consecutive recordings of 100 and 300 traces. The waveform’s initial positive peak, designated as P1, typically occurred around 100 ms, followed by a second negative peak, referred to as N2. To assess the impact of injury, the mean amplitude of the P1-N2 wave in the affected eye was compared to that of the contralateral control eye, yielding a percentage change in amplitude. The investigators responsible for amplitude measurement were blinded to the treatment conditions to ensure objectivity.

### Optokinetic response (OKR) measurement

Mouse spatial visual acuity was assessed using the OKR assay, as previously described^19,20^. Unrestrained live mice were placed on a central platform surrounded by four 17-inch LCD monitors (Dell, Phoenix, AZ). Their movements were recorded with an overhead video camera. A virtual rotating cylinder with vertical sine wave gratings was presented on the monitors using OptoMotry software. (Cerebral Mechanics Inc., Lethbridge, Alberta, Canada). The rotation of the grating created a virtual reality environment to evaluate spatial acuity, with clockwise rotation stimulating the left eye and counterclockwise rotation stimulating the right eye. Once the mice became stationary, the visual stimulus was initiated at a low spatial frequency of 0.1 cycles/degree for a duration of five seconds. During this interval, the investigator observed whether the mouse exhibited head tracking movements in response to the rotating grating. The brief exposure time was designed to minimize adaptation to the stimulus, thereby reducing the likelihood of false-negative results. Mice that demonstrated head tracking were considered capable of perceiving the grating. The spatial frequency was then incrementally increased until the highest frequency that the mouse could reliably track was identified. Relative spatial vision was determined by calculating the ratio of the maximum spatial frequency detected by the treated eye to that of the contralateral control eye, expressed as a percentage. To ensure consistency in experimental conditions, all OKR assessments were conducted during the morning hours. The investigators performing the measurements were blinded to the treatment groups to maintain objectivity and eliminate potential bias in the analysis.

### Optic nerve clearance and axon regeneration quantification

Optic nerves traced anterogradely with cholera toxin subunit B (CTB) were prepared for visualization and analysis using an adapted version of the iDISCO clearing protocol (17). Intravitreal injection of CTB-555 was conducted 48 hours prior to the perfusion of the animals with 4% paraformaldehyde (PFA) in phosphate-buffered saline (PBS). Initially, the optic nerves were rinsed in phosphate-buffered saline (PBS) for 30 minutes, followed by sequential dehydration in a graded methanol series (20%, 40%, 60%, 80%, and 100% in PBS), with each step lasting 30 minutes. The samples were then incubated in a dichloromethane (DCM)/methanol solution (2:1) for 30 minutes, followed by two consecutive 30-minute immersions in 100% DCM and dibenzyl ether (DBE), respectively. Cleared optic nerves were subsequently mounted between two 22 × 22 mm coverslips in DBE, sealed with transparent nail polish, and prepared for imaging.

Imaging was performed using a 25× oil immersion objective in airy scan mode, acquiring Z-stacks at 6 µm intervals along the length of the optic nerve. To capture the entire nerve, tile scanning was employed. Axon quantification was conducted as previously described^12^. Briefly, axons labeled with CTB were counted at specified distances from the crush site (e.g., 250 µm, 500 µm, 1000 µm, 1500 µm, and every 250 µm thereafter) by assessing the number of fibers crossing perpendicular lines drawn on optical sections of the nerve. To estimate axon density, Z-stacks were sampled at three depths (60, 120, and 180 µm), and the number of axons was calculated using the formula: Axon Number=(Counted Axons)/(R*t) where (R) represents the width of the stack at the counting location, and (t) is the optical section thickness (6 µm). The total axon number was calculated using mean axon density from these three stacks using the formula:∑ad = πr^2^ * mean axon density where (r) is the optic nerve radius. Only CTB signals within a predefined intensity threshold (established post-background subtraction) were considered as individual axons. All axon counts were performed by investigators who were blinded to the treatment groups to ensure unbiased results.

### Immunohistochemistry and quantification

The detailed procedures have been published previously^12,30,31^. Following transcardiac perfusion with 4% paraformaldehyde (PFA) in PBS, mice eyes were carefully enucleated and post-fixed in 4% PFA at room temperature for 2 hours. The tissues were subsequently cryoprotected by incubation in 30% sucrose overnight at 4°C. Retinas were then isolated, rinsed thoroughly in PBS, and subjected to blocking in a staining buffer containing 10% normal goat serum and 2% Triton X-100 in PBS for 30 minutes at room temperature. To visualize retinal ganglion cells (RGCs) and 3-HA expression, retinas were incubated with primary antibodies at 4°C overnight. Specifically, an anti-RBPMS guinea pig antibody (ProSci, California) was applied at a dilution of 1:4000, while a rat anti-3HA antibody was used at a dilution of 1:500. Following primary antibody incubation, retinas underwent three washes with PBS (30 minutes per wash). Secondary antibody incubation was then performed at room temperature for 1 hour using Alexa Fluor 647-conjugated goat anti-guinea pig, Alexa Fluor 488-conjugated goat anti-guinea pig, and Cy3 or Cy5-conjugated goat anti-rat antibodies (1:200 dilution; Jackson ImmunoResearch, West Grove, Pennsylvania). Retinas were washed three additional times with PBS (30 minutes per wash) before mounting with Fluoromount-G (Southern Biotech, Birmingham, Alabama) under a coverslip. Immunostained retinal whole mounts were imaged using a Keyence fluorescence microscope and a Zeiss LSM 880 confocal microscope equipped with 20× and 40× oil immersion objectives. For RGC quantification, 6–9 fields of view (each measuring 332 × 332 µm) were sampled from the peripheral, middle, and central regions of each retina. Imaging and stitching were conducted using Keyence fluorescence microscope (BZ-X800, Itasca), and RBPMS-positive RGCs were quantified using Fiji/ImageJ software. RGC survival (%) was determined by comparing the number of surviving RGCs in the diseased eye with that in the contralateral uninjured eye. All cell counts were performed by investigators blinded to the experimental conditions to ensure unbiased analysis.

For cross-sectional analysis, mouse eyeballs were embedded in Tissue-Tek OCT compound and rapidly frozen on dry ice, facilitating subsequent cryosectioning with a Leica cryostat. The primary antibodies employed for immunostaining included anti-RBPMS (1:4,000; custom-made by ProSci), anti-4-Hydroxynonenal (1:200; catalog number MAB3249-SP, Novus), and anti-Glutathione Peroxidase 4 (1:200; catalog number ab125066, Abcam). Following the application of primary antibodies, sections were incubated with secondary antibodies at room temperature for 1 hour. The secondary antibodies used included Alexa Fluor 594- or Alexa Fluor 647-conjugated goat anti-rabbit, Alexa Fluor 488-conjugated goat anti-guinea pig, and Alexa Fluor 594-conjugated goat anti-mouse antibodies, all at a dilution of 1:200 (Jackson ImmunoResearch, West Grove, Pennsylvania). For quantification of relative fluorescence intensity, three immunofluorescence images of the retina were captured for each sample across all experimental groups using a Keyence fluorescence microscope (BZ-X800, Itasca), with triplicate samples in independent experiments. The immunofluorescence intensity of the ganglion cell layer (GCL) was analyzed using Fiji/ImageJ software, following background subtraction. The final relative intensity was calculated using the formula: (Mean immunofluorescence intensity — Background immunofluorescence intensity) / Background immunofluorescence intensity.

### Preparation and Quantification of Optic Nerve Semi-Thin Sections

Optic nerve semi-thin sectioning and axon survival analysis were conducted following a previously established protocol. Briefly, transverse optic nerve sections of 1 µm thickness were prepared using an ultramicrotome (EM UC7, Leica, Wetzlar, Germany). Tissue samples were collected 2 mm distal to the globe and approximately 1.5 mm from the crush site. Sections were stained with 1% para-phenylenediamine (PPD) dissolved in a 1:1 methanol mixture to enhance contrast. Entire optic nerves were imaged using a Keyence fluorescence microscope equipped with a 100× objective lens. High-resolution images were acquired and stitched together to generate a comprehensive representation of the nerve. Axon survival was quantified by applying a grid of automated 10 µm × 10 µm counting frames to approximately 10% of the total nerve area using the “AxonCounter” plugin in ImageJ^26^. Within the defined grid, axons were manually identified and enumerated using the ImageJ cell counter tool. For each optic nerve, the mean axon count across all analysed images was determined. The percentage of surviving axons was calculated by normalizing the mean axon count of the injured nerve to that of the contralateral uninjured control. To ensure unbiased quantification, investigators performing axon analysis were blinded to the experimental conditions.

### Statistical analyses

All statistical analyses and data visualization were performed using GraphPad Prism (version 10) or R package. Data are presented as mean ± standard error of the mean (SEM). Comparisons between two groups were conducted using an unpaired Student’s *t*-test. For multiple group comparisons, one-way ANOVA followed by an appropriate post hoc test or two-way ANOVA was applied, depending on the experimental design. Statistical significance was defined as *p* < 0.05.

## Acknowledgement

We thank the Hu lab members for critical discussion. We are grateful for an unrestricted grant from Research to Prevent Blindness and NEI P30 EY026877 to the Department of Ophthalmology. M.Y. is supported by 2024 ARVO Foundation Early Career Clinician-Scientist Research Award. Y.H. is supported by NIH/NEI grants EY032518, EY024932, EY023295, EY034353, and grants from Glaucoma Research Foundation (CFC3), Chan Zuckerberg Initiative NDCN Collaborative Pairs Projects, Stanford SPARK program, Stanford Innovative Medicines Accelerator, and RPB Stein Innovation Award.

## Author Contributions

M.Y. and Y.H., designed the experiments. M.Y., D. L., F.B., X.F., L. Li, H.H. established methodology and M.Y., F.B., D. Liu., L. Li, R. D., F.C., P.O., and A.L. involved in sample collection and quantification. L. Liu. produced AAVs. M.Y. and Y.H. prepared the manuscript with support from all the authors.

---

**Figure S1.**
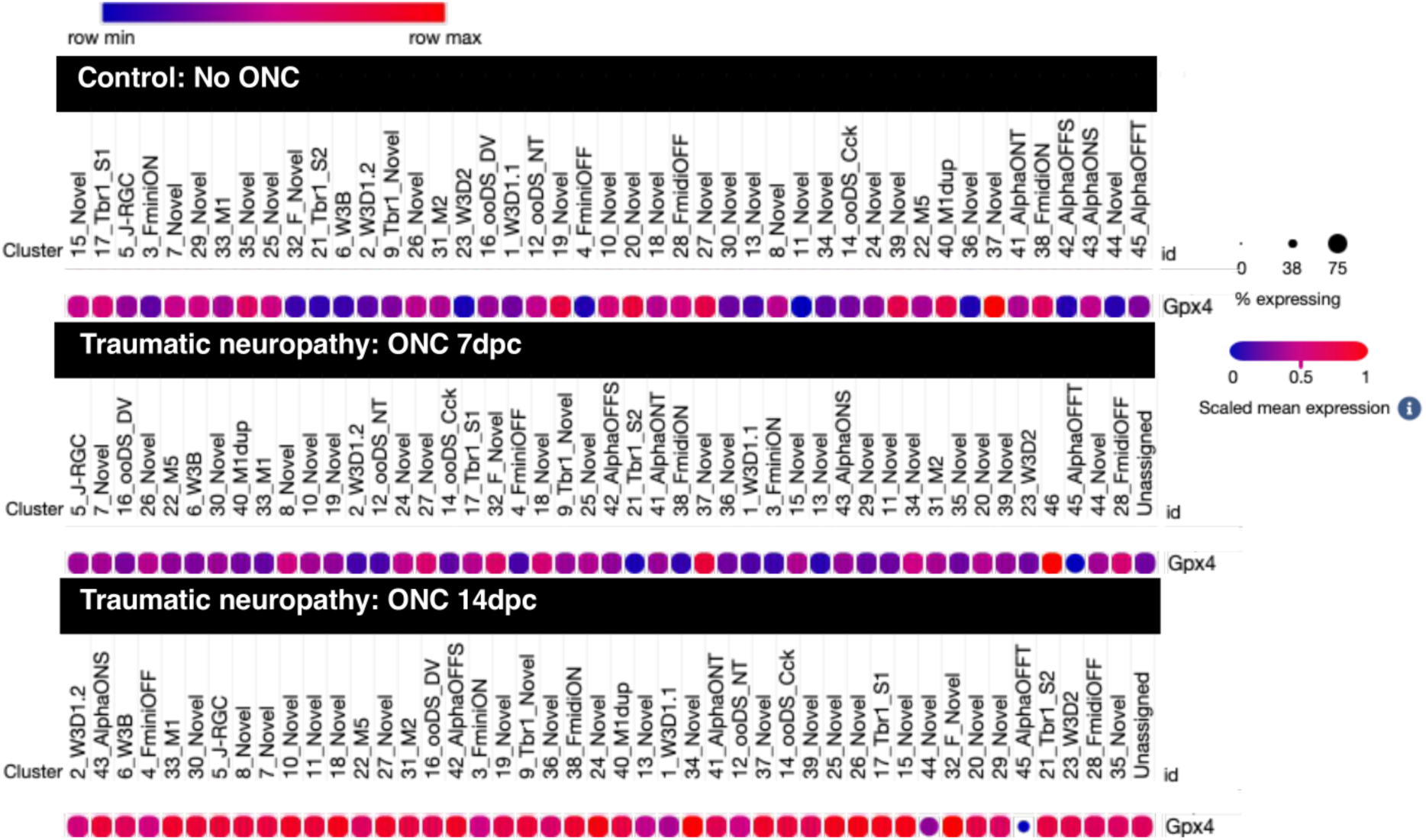
The transcriptome profiles of GPX4 in RGC subtypes after ONC. Data was obtained from online source: https://singlecell.broadinstitute.org/single_cell/study/SCP509/mouse-retinal-ganglion-cell-adult-atlas-and-optic-nerve-crush-time-series#study-visualize). The mRNA levels of GPX4 in individual RGC subtypes were identified by scRNA-seq analysis^11^. Data are presented as dot plots. The dot size shows % of expression, and the dot color shows scaled mean expression.

**Figure S2.**
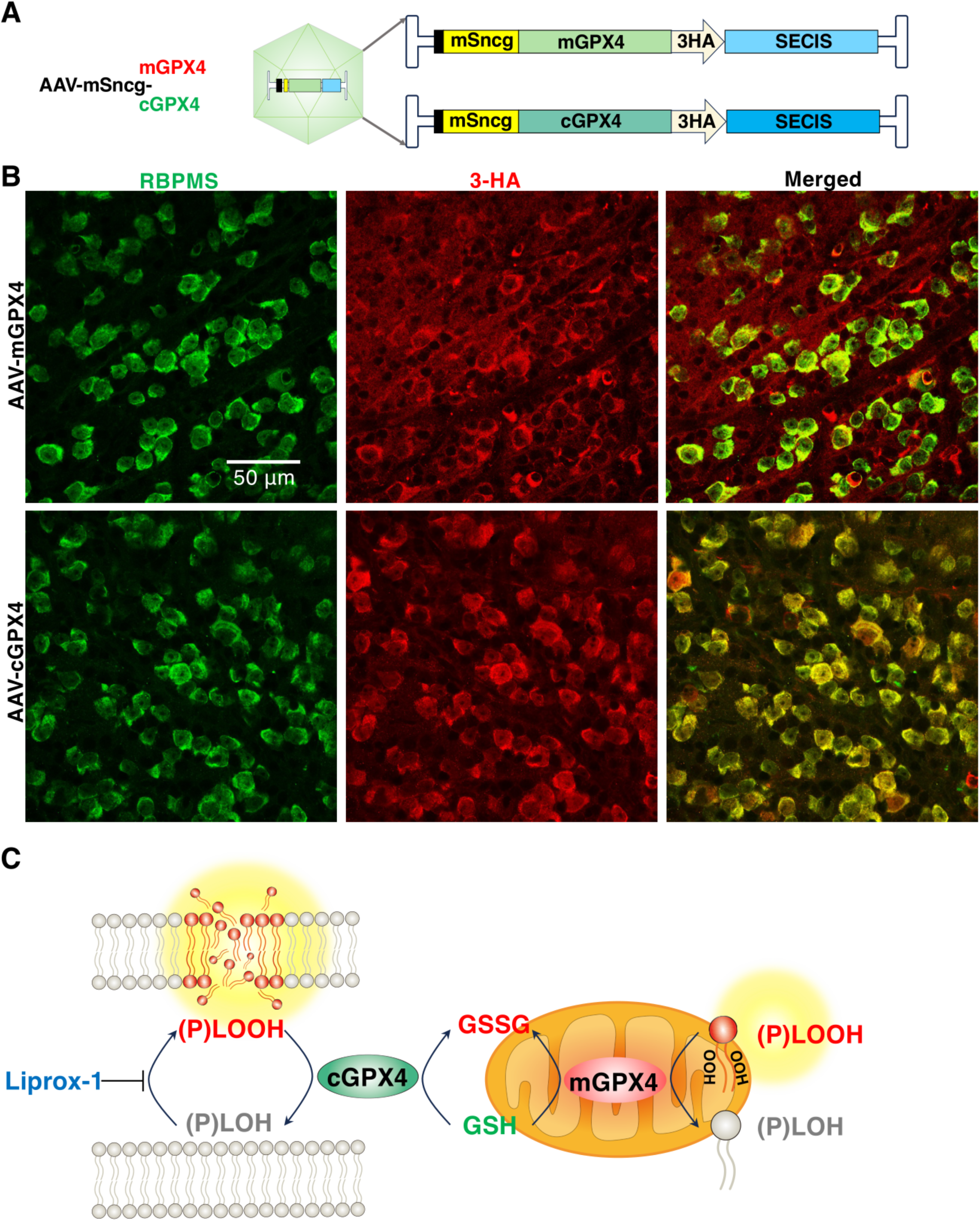
RGC-specific expression of cGPX4 and mGPX4. (**A**) AAV vectors of cGPX4 and mGPX4 with 3HA tag and SECIS (Selenocysteine Insertion Sequence), under mSncg promoter. (**B**) AAV-mediated transgene expression in RGCs labelled by HA antibodies 2 weeks after AAV intravitreal injection. Scale bar, 50µm. (**C**) Scheme of the function of GPX4 and Liprox-1 in the inhibition of lipid peroxidation.

**Figure S3.**
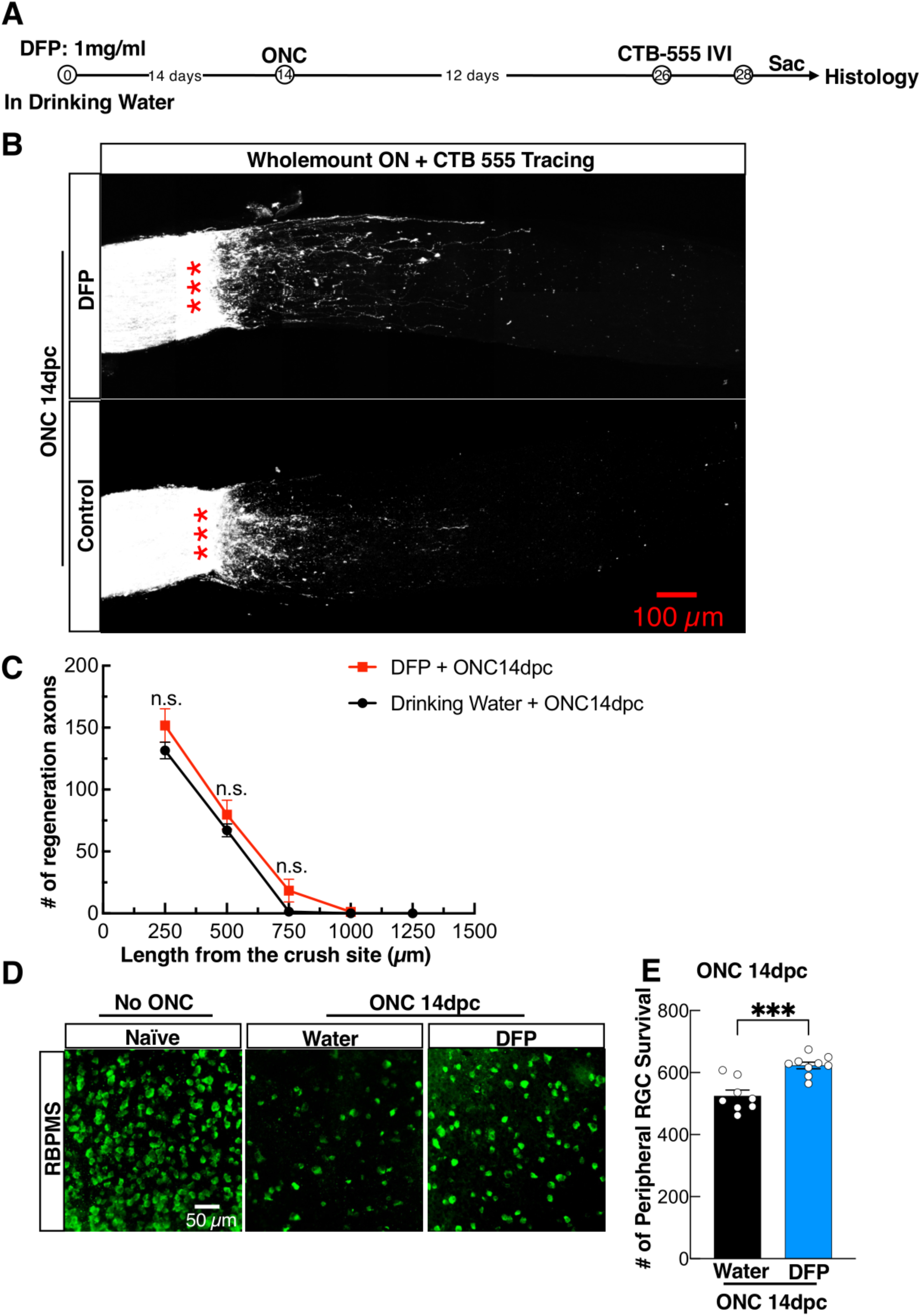
Iron chelator DFP did not promote ON regeneration but promoted significant RGC survival after ONC. (**A**) A flowchart showing the experimental design. (**B**) Confocal images of ON wholemount after optical clearance showing maximum intensity projection of regenerating fibers labelled with CTB-Alexa 555 at 14dpc. Scale bar, 100 μm. ***: Crush site. (**C**) Quantification of regenerating fibers at different distances distal to the lesion site. Data are presented as means ± s.e.m, n=8 of DFP group, n=11 of vehicle control group. Data are presented as means ± s.e.m, n.s.: no statistical significance. Two-way ANOVA with Tukey’s multiple comparisons test. (D). Representative confocal images of the flat-mounted retinas at the peripheral regions showing surviving RBPMS-positive RGCs at 14dpc. Scale bar, 50 µm. (E). Quantification of surviving RGC somata from peripheral flat-mounted retinas, DFP (n=9), Water (n=8). Data are presented as means ± s.e.m, ***: p < 0.001, Student’s t-test.

**Figure S4.**
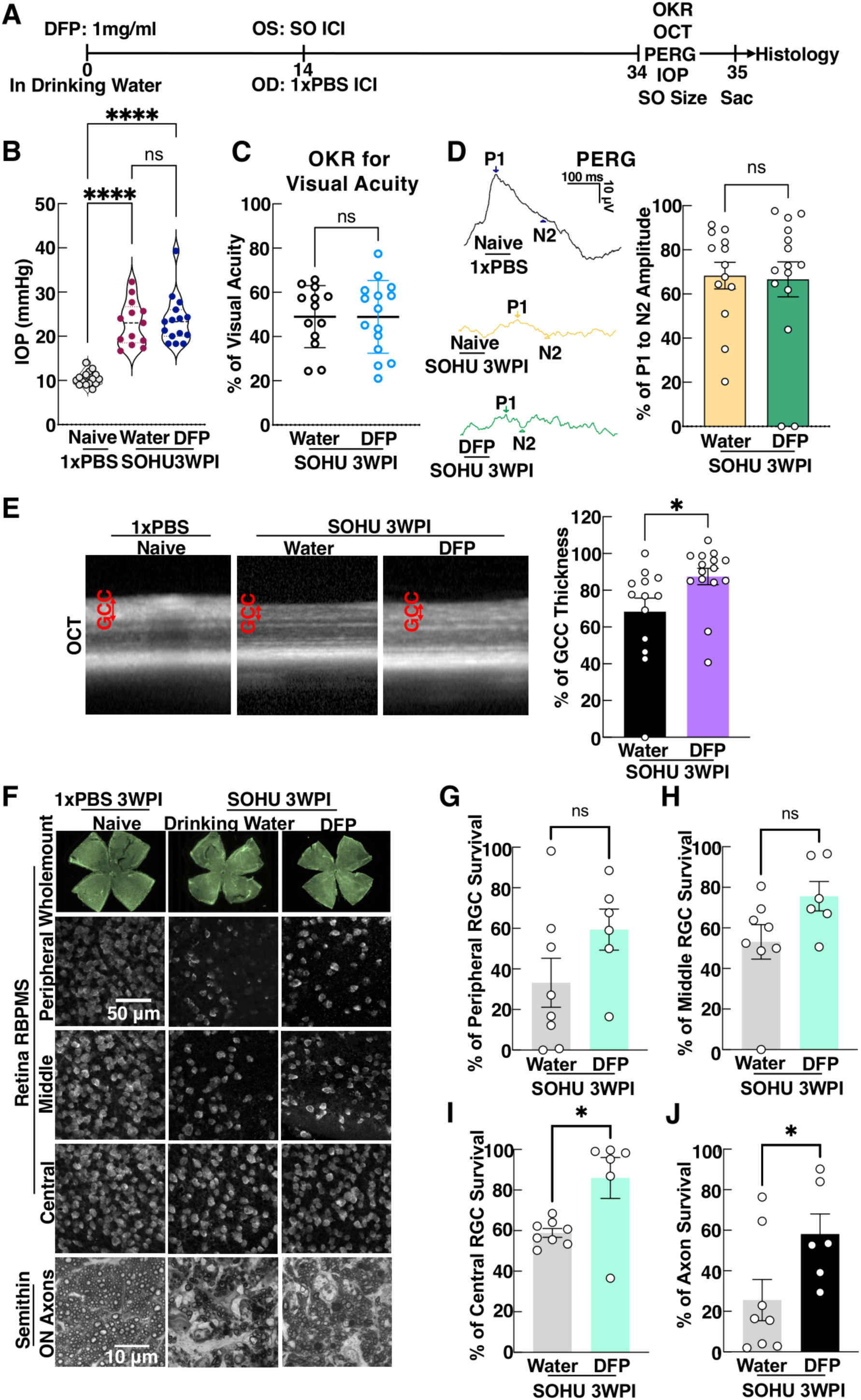
Iron chelator DFP did not promote significant neuroprotection in the SOHU glaucoma model. (**A**) Timeline of DFP treatment daily starting 14 days before SO intracameral injection until the termination of the experiments. OS: the left eye, OD: the right eye, ICL: intracameral injection, SO: silicon oil, Sac: Sacrificed. (**B**) IOP measurement at 3WPI. Naive mice with intracameral injection of 1xPBS as naïve control (n=15 mice), vehicle (drinking water)-treated SOHU mice (n=13 mice), and DFP-treated SOHU mice (n=15 mice). Data are presented as means ± s.e.m. ns: no statistical significance, ****: *p* < 0.0001, one-way ANOVA with Tukey’s multiple comparison tests. (**C**) Visual acuity measurement by optomotor, represented as the percentage of glaucomatous eyes compared to the contralateral control eyes. (**D**) Left, representative waveforms of PERG. Right, quantification of P1-N2 amplitude of PERG at 3WPI, represented as a percentage of glaucomatous eyes compared to the contralateral control eyes. (**E**) Left: Representative *in vivo* OCT images of mouse retinas at 3WPI. GCC: including RNFL, GCL, and IPL layers; indicated as double-end arrows. Right: Quantification of GCC thickness was measured by OCT at 3WPI, represented as the percentage of GCC thickness in the SOHU eyes compared to the contralateral control eyes. (**F**) The top panel: representative images of the whole flat-mounted retinas showing surviving RBPMS-positive RGCs at 3WPI. The middle 3 panels: representative images of peripheral, middle, and central retinas; scale bar: 50µm. The bottom panel: representative confocal microscope images of ON semi-thin transverse sections with paraphenylenediamine staining at 3WPI. Scale bar: 10µm. (**G,H,I**) Quantification of surviving RBPMS+ RGCs at peripheral, middle, and central retinas, represented as a percentage of glaucomatous eyes compared to the contralateral control eyes. vehicle control group (n=8 mice), DFP-treated group (n=6 mice). (**J**) Quantification of surviving axons in ON semithin sections, represented as the percentage of glaucomatous eyes compared with the contralateral control eyes. vehicle control group (n=8 mice), DFP-treated group (n=6 mice). All the quantification data of C,D,G-J are presented as means ± s.e.m, ns, no statistical significance, *: p < 0.05, Student’s t-test.

## References

1. Tham, Y.C., Li, X., Wong, T.Y., Quigley, H.A., Aung, T., and Cheng, C.Y. (2014). Global prevalence of glaucoma and projections of glaucoma burden through 2040: a systematic review and meta-analysis. Ophthalmology 121, 2081–2090. 10.1016/j.ophtha.2014.05.013.

2. Quigley, H.A., Nickells, R.W., Kerrigan, L.A., Pease, M.E., Thibault, D.J., and Zack, D.J. (1995). Retinal ganglion cell death in experimental glaucoma and after axotomy occurs by apoptosis. Invest Ophthalmol Vis Sci 36, 774–786.

3. Nickells, R.W., Howell, G.R., Soto, I., and John, S.W. (2012). Under pressure: cellular and molecular responses during glaucoma, a common neurodegeneration with axonopathy. Annu Rev Neurosci 35, 153–179. 10.1146/annurev.neuro.051508.135728.

4. Calkins, D.J. (2021). Adaptive responses to neurodegenerative stress in glaucoma. Prog Retin Eye Res 84, 100953. 10.1016/j.preteyeres.2021.100953.

5. Guo, M., Zhu, Y., Shi, Y., Meng, X., Dong, X., Zhang, H., Wang, X., Du, M., and Yan, H. (2022). Inhibition of ferroptosis promotes retina ganglion cell survival in experimental optic neuropathies. Redox Biol 58, 102541. 10.1016/j.redox.2022.102541.

6. Yao, F., Peng, J., Zhang, E., Ji, D., Gao, Z., Tang, Y., Yao, X., and Xia, X. (2023). Pathologically high intraocular pressure disturbs normal iron homeostasis and leads to retinal ganglion cell ferroptosis in glaucoma. Cell Death Differ 30, 69–81. 10.1038/s41418-022-01046-4.

7. Ye, H., Feng, Y., Xiang, W., Lin, Z., Li, Y., Hu, W., Liu, K., Tao, S., Shu, Q., Wang, J., Xu, F., et al. (2025). Ferroptosis Contributes to Retinal Ganglion Cell Loss in GLAST Knockout Mouse Model of Normal Tension Glaucoma. Invest Ophthalmol Vis Sci 66, 26. 10.1167/iovs.66.5.26.

8. Dixon, S.J., Lemberg, K.M., Lamprecht, M.R., Skouta, R., Zaitsev, E.M., Gleason, C.E., Patel, D.N., Bauer, A.J., Cantley, A.M., Yang, W.S., Morrison, B., 3rd, et al. (2012). Ferroptosis: an iron-dependent form of nonapoptotic cell death. Cell 149, 1060-1072. 10.1016/j.cell.2012.03.042.

9. Jiang, X., Stockwell, B.R., and Conrad, M. (2021). Ferroptosis: mechanisms, biology and role in disease. Nat Rev Mol Cell Biol 22, 266–282. 10.1038/s41580-020-00324-8.

10. Friedmann Angeli, J.P., Schneider, M., Proneth, B., Tyurina, Y.Y., Tyurin, V.A., Hammond, V.J., Herbach, N., Aichler, M., Walch, A., Eggenhofer, E., Basavarajappa, D., et al. (2014). Inactivation of the ferroptosis regulator Gpx4 triggers acute renal failure in mice. Nat Cell Biol 16, 1180–1191. 10.1038/ncb3064.

11. Tran, N.M., Shekhar, K., Whitney, I.E., Jacobi, A., Benhar, I., Hong, G., Yan, W., Adiconis, X., Arnold, M.E., Lee, J.M., Levin, J.Z., et al. (2019). Single-Cell Profiles of Retinal Ganglion Cells Differing in Resilience to Injury Reveal Neuroprotective Genes. Neuron 104, 1039–1055 e1012. 10.1016/j.neuron.2019.11.006.

12. Li, L., Fang, F., Feng, X., Zhuang, P., Huang, H., Liu, P., Liu, L., Xu, A.Z., Qi, L.S., Cong, L., and Hu, Y. (2022). Single-cell transcriptome analysis of regenerating RGCs reveals potent glaucoma neural repair genes. Neuron 110, 2646–2663 e2646. 10.1016/j.neuron.2022.06.022.

13. Arai, M., Imai, H., Sumi, D., Imanaka, T., Takano, T., Chiba, N., and Nakagawa, Y. (1996). Import into mitochondria of phospholipid hydroperoxide glutathione peroxidase requires a leader sequence. Biochem Biophys Res Commun 227, 433–439. 10.1006/bbrc.1996.1525.

14. Wang, Q., Zhuang, P., Huang, H., Li, L., Liu, L., Webber, H.C., Dalal, R., Siew, L., Fligor, C.M., Chang, K.C., Nahmou, M., et al. (2020). Mouse gamma-Synuclein Promoter-Mediated Gene Expression and Editing in Mammalian Retinal Ganglion Cells. J Neurosci 40, 3896–3914. 10.1523/JNEUROSCI.0102-20.2020.

15. Zilka, O., Shah, R., Li, B., Friedmann Angeli, J.P., Griesser, M., Conrad, M., and Pratt, D.A. (2017). On the Mechanism of Cytoprotection by Ferrostatin-1 and Liproxstatin-1 and the Role of Lipid Peroxidation in Ferroptotic Cell Death. ACS Cent Sci 3, 232–243. 10.1021/acscentsci.7b00028.

16. Devisscher, L., Van Coillie, S., Hofmans, S., Van Rompaey, D., Goossens, K., Meul, E., Maes, L., De Winter, H., Van Der Veken, P., Vandenabeele, P., Berghe, T.V., et al. (2018). Discovery of Novel, Drug-Like Ferroptosis Inhibitors with in Vivo Efficacy. J Med Chem 61, 10126–10140. 10.1021/acs.jmedchem.8b01299.

17. Gawel, S., Wardas, M., Niedworok, E., and Wardas, P. (2004). [Malondialdehyde (MDA) as a lipid peroxidation marker]. Wiad Lek 57, 453–455.

18. Zarkovic, N. (2003). 4-hydroxynonenal as a bioactive marker of pathophysiological processes. Mol Aspects Med 24, 281–291. 10.1016/s0098-2997(03)00023-2.

19. Fang, F., Zhang, J., Zhuang, P., Liu, P., Li, L., Huang, H., Webber, H.C., Xu, Y., Liu, L., Dalal, R., Sun, Y., et al. (2021). Chronic mild and acute severe glaucomatous neurodegeneration derived from silicone oil-induced ocular hypertension. Scientific reports 11, 9052. 10.1038/s41598-021-88690-x.

20. Zhang, J., Li, L., Huang, H., Fang, F., Webber, H.C., Zhuang, P., Liu, L., Dalal, R., Tang, P.H., Mahajan, V.B., Sun, Y., et al. (2019). Silicone oil-induced ocular hypertension and glaucomatous neurodegeneration in mouse. eLife 8. 10.7554/eLife.45881.

21. Zhang, J., Fang, F., Li, L., Huang, H., Webber, H.C., Sun, Y., Mahajan, V.B., and Hu, Y. (2019). A Reversible Silicon Oil-Induced Ocular Hypertension Model in Mice. Journal of visualized experiments : JoVE 153. 10.3791/60409.

22. Moshiri, A., Fang, F., Zhuang, P., Huang, H., Feng, X., Li, L., Dalal, R., and Hu, Y. (2022). Silicone Oil-Induced Glaucomatous Neurodegeneration in Rhesus Macaques. Int J Mol Sci 23. 10.3390/ijms232415896.

23. Benowitz, L.I., He, Z., and Goldberg, J.L. (2017). Reaching the brain: Advances in optic nerve regeneration. Exp Neurol 287, 365–373. 10.1016/j.expneurol.2015.12.015.

24. Paez-Perez, M., Vysniauskas, A., Lopez-Duarte, I., Lafarge, E.J., Lopez-Rios De Castro, R., Marques, C.M., Schroder, A.P., Muller, P., Lorenz, C.D., Brooks, N.J., and Kuimova, M.K. (2023). Directly imaging emergence of phase separation in peroxidized lipid membranes. Commun Chem 6, 15. 10.1038/s42004-022-00809-x.

25. Balakrishnan, M., and Kenworthy, A.K. (2024). Lipid Peroxidation Drives Liquid-Liquid Phase Separation and Disrupts Raft Protein Partitioning in Biological Membranes. J Am Chem Soc 146, 1374–1387. 10.1021/jacs.3c10132.

26. Fang, F., Liu, P., Huang, H., Feng, X., Li, L., Sun, Y., Kaufman, R.J., and Hu, Y. (2023). RGC-specific ATF4 and/or CHOP deletion rescues glaucomatous neurodegeneration and visual function. Mol Ther Nucleic Acids 33, 286–295. 10.1016/j.omtn.2023.07.015.

27. Fang, F., Zhuang, P., Feng, X., Liu, P., Liu, D., Huang, H., Li, L., Chen, W., Liu, L., Sun, Y., Jiang, H., et al. (2022). NMNAT2 is downregulated in glaucomatous RGCs, and RGC-specific gene therapy rescues neurodegeneration and visual function. Mol Ther 30, 1421–1431. 10.1016/j.ymthe.2022.01.035.

28. Zhang, J., Fang, F., Li, L., Huang, H., Webber, H.C., Sun, Y., Mahajan, V.B., and Hu, Y. (2019). A Reversible Silicon Oil-Induced Ocular Hypertension Model in Mice. J Vis Exp. 10.3791/60409.

29. Li, L., Huang, H., Fang, F., Liu, L., Sun, Y., and Hu, Y. (2020). Longitudinal Morphological and Functional Assessment of RGC Neurodegeneration After Optic Nerve Crush in Mouse. Frontiers in cellular neuroscience 14, 109. 10.3389/fncel.2020.00109.

30. Fang, F., Zhuang, P., Feng, X., Liu, P.T., Liu, D., Huang, H.L., Li, L., Chen, W., Liu, L., Sun, Y., Jiang, H.W., et al. (2022). NMNAT2 is downregulated in glaucomatous RGCs, and RGC-specific gene therapy rescues neurodegeneration and visual function. Molecular Therapy 30, 1421–1431. 10.1016/j.ymthe.2022.01.035.

31. Liu, D., Webber, H.C., Bian, F., Xu, Y., Prakash, M., Feng, X., Yang, M., Yang, H., You, I.J., Li, L., Liu, L., et al. (2025). Optineurin-facilitated axonal mitochondria delivery promotes neuroprotection and axon regeneration. Nat Commun 16, 1789. 10.1038/s41467-025-57135-8.

